# Convergent molecular changes are associated with the evolution of low susceptibility to the ash dieback pathogen

**DOI:** 10.1101/2025.02.08.637240

**Authors:** Laura J. Kelly, Sarah E. R. Coates, Steve J. Lee, Stephen J. Rossiter, Richard J. A. Buggs

**Author notes:** Author for correspondence: Laura J. Kelly, Royal Botanic Gardens, Kew, Richmond, Surrey, UK, Richard J. A. Buggs, Royal Botanic Gardens, Kew, Richmond, Surrey, UK. Molecular Ecology and Evolution group, School of Environmental and Natural Sciences, Bangor University, Bangor, UK; Royal Botanic Gardens, Kew, Richmond, Surrey, UK.

## Abstract

Non-native pests and pathogens increasingly threaten global forest ecosystems. An understanding of the genomic basis of low susceptibility in the natural hosts of such pests and pathogens, with which they share a coevolutionary history, can enhance restoration efforts by facilitating the selection of individuals carrying beneficial alleles. Within the genus *Fraxinus* (ash trees) low susceptibility to the fungal pathogen *Hymenoscyphus fraxineus*, the causative agent of the ash dieback disease (ADB) epidemic, is observed in three independent lineages of known or plausible natural hosts. Here, we seek to elucidate the genetic basis of this trait, which is key to the future survival of *Fraxinus excelsior* populations in Europe, using a molecular convergence approach. We analysed 4,300 protein-coding loci for amino acid convergence between lineages with low susceptibility. After filtering for potential false positives, we find 62 genes that have a signal of excess convergence between *Fraxinus* lineages with low ADB susceptibility. Eleven of these loci have additional evidence for a role in defence against fungal pathogens, with a further 17 linked to more general immunity or defence, or other functions relevant to the response against ADB such as cell wall biogenesis. The candidate loci discovered here complement those reported by previous genomic studies of *F. excelsior*, and can be targeted in efforts to mitigate the devastation caused by this deadly disease by informing breeding programmes involving hybridisation and marker-assisted back-crossing. Our study demonstrates the benefit of genomic analyses incorporating natural hosts, when seeking to tackle biotic threats to naïve host populations.

**Significance Statement:** Knowledge of the genomic basis of variation in susceptibility to emerging forest pests and pathogens can provide the foundation for interventions to mitigate such threats, but studies restricted to the analysis of naïve host populations may miss relevant genomic loci that could be present in coevolved hosts. Using comparative genomic analyses for the detection of molecular convergence between lineages of ash trees with low susceptibility to ash dieback, incorporating known and plausible natural hosts, we detect novel candidate loci for defence against this disease. Our findings can be used to support efforts to tackle one of the world’s worst forest pathogens and demonstrate the value of integrating genomic data from coevolved hosts when aiming to identify the basis of low susceptibility to biotic threats.

## Introduction

The identification of genomic variants underlying trait variation is a key aim in biology, helping to address long standing evolutionary questions (Smith et al. 2020), as well as having important practical applications (e.g. (Kersey et al. 2020; Theissinger et al. 2023)). Where the trait in question is the ability to survive a destructive pest or pathogen, such knowledge can provide the basis for interventions to enhance the future viability of affected species (Theissinger et al. 2023). Tree health epidemics caused by non-native insects and diseases represent one of the most critical threats to global forest ecosystems (Williams et al. 2023). Invasive pathogens such as *Cryphonectria parasitica*, the causal agent of chestnut blight, can bring about continent-scale impacts and functional extinction of keystone tree species (Williams et al. 2023). Although naïve host populations may have partial resistance to these threats through exaptation of standing genetic variation reflecting selection by other stressors (Budde et al. 2016) and have the evolutionary potential to adapt to novel pests or pathogens so that a stable coexistence is established (Budde et al. 2016; Ennos 2015), safeguarding vulnerable tree species may also require human interventions, such as resistance breeding programmes (Ennos 2015). In cases where the natural hosts of pests and pathogens are known, these can provide a valuable source of adaptive genetic variation that reflects the coevolutionary history of these organisms, which could supplement that found within naïve hosts.

Ash dieback (ADB), caused by the non-native invasive fungal pathogen *Hymenoscyphus fraxineus* (Kowalski 2006), is a recently emerging infectious disease with devastating impacts on ash trees within Europe. ADB was first reported in Poland in the early 1990s and over the following decades has caused high levels of mortality in *Fraxinus excelsior* (European ash) populations throughout the continent (Enderle et al. 2019; Marçais et al. 2022; Coker et al. 2019), also putting at risk dozens of other species that use ash as a host (Mitchell et al. 2014; Hultberg et al. 2020). Another of the three *Fraxinus* species native to Europe, *F. angustifolia* (narrow-leaved ash), is also highly susceptible to the disease, although populations in the more southerly part of its range are unaffected thus far (Marçais et al. 2022). A small proportion of *F. excelsior* and *F. angustifolia* individuals have low susceptibility to ADB, with most estimates indicating this applies to less than 10% of trees (Enderle et al. 2019; Schertler et al. 2026), meaning they develop no or only minor disease symptoms. Moreover, heritability estimates indicate a significant proportion of the variance in susceptibility can be explained by genetic differences (McKinney et al. 2014; Hauptman et al. 2016) and genomic variants contributing to this polygenic trait have been identified in *F. excelsior* (Stocks et al. 2019; Doonan et al. 2025). Although there is already evidence for an adaptive response that could increase the frequency of individuals with low susceptibility (Metheringham et al. 2025), it may be beneficial to accelerate this process via selective breeding programmes (Evans 2019; Plumb et al. 2020; Metheringham et al. 2025; Schertler et al. 2026). Such programmes have been established in several European countries (e.g. Langer et al. (2022)) and could be further enhanced by knowledge of genetic variants associated with low susceptibility to ADB in natural, coevolved, hosts of *H. fraxineus*, which may be absent from species such as *F. excelsior*.

To date, two natural host species have been recorded for *H. fraxineus* within its native range in East Asia and the Russian Far East (Baral & Bemmann 2014): *F. mandshurica* and *F. chinensis* subsp. *rhynchophylla* (Cleary et al. 2016; Drenkhan, Solheim, et al. 2017; Zheng & Zhuang 2014; Han et al. 2014; Zhao et al. 2013). In these hosts, *H. fraxineus* occurs asymptomatically as a leaf endophyte (Cleary et al. 2016; Inoue et al. 2019), or a pathogen causing minor disease symptoms (Drenkhan, Solheim, et al. 2017), but the severe dieback and mortality observed in Europe have not been reported in their native range (Drenkhan, Solheim, et al. 2017; Cleary et al. 2016; Han et al. 2014; Zheng & Zhuang 2014). East Asian strains of *H. fraxineus* are as virulent as those found in Europe (Gross & Sieber 2016; Gross & Holdenrieder 2015) suggesting the absence of serious disease within the native range may be due to lower susceptibility of hosts that have coevolved with the fungus. In addition to these confirmed natural hosts, susceptibility of numerous other *Fraxinus* species has been assessed, either from natural infections within the introduced range of the fungus or experimentally. These reveal the lowest susceptibility is found among ash species native to East Asia (Nielsen et al. 2017; Drenkhan, Agan, et al. 2017; Pastirčáková et al. 2020; Gross & Sieber 2016; Queloz et al. 2017; Hiemstra et al. 2019). Although the full extent of the native distribution of *H. fraxineus* is unknown, it has been recorded in China, Japan, Republic of Korea and the Russian Far East (Marçais et al. 2022). Thus, further ash species native to these regions may have evolved defence responses to the fungus and could provide a source of genomic variants for use in efforts to enhance the resilience of European native ash species.

Here we use existing genomic data from *Fraxinus* (Kelly et al. 2020) to analyse multiple independent lineages with contrasting levels of susceptibility to ADB within the genus, with the aim of detecting variants contributing to low susceptibility to the pathogen. We take an approach that involves the detection of molecular convergence (or parallel genetic changes - we do not attempt here to make a distinction between these (Elmer & Meyer 2011)), one of a group of phylogenetic genotype-to-phenotype methods that can be effective for identification of candidate loci (Macdonald et al. 2025), as the repeated evolution of phenotypic traits may involve reuse of the same genes or genetic variants (Stern 2013; Storz 2016). Since ash species with low susceptibility to ADB span three separate clades within the genus (Kelly et al. 2020) this trait may have evolved (or been preserved from a common ancestor) on multiple occasions, making it possible to use such an approach to identify genomic variants associated with this trait variation (Smith et al. 2020). We tested thousands of nuclear genes for evidence of molecular convergence at the amino acid level associated with low susceptibility to this deadly disease. We uncover dozens of genes with potential roles in defence against *H. fraxineus* in its natural hosts, most of which have not been identified by genome-wide analyses within *F. excelsior*, demonstrating the potential to identify genetic variation associated with key adaptive traits from analyses of coevolved hosts.

## Results

### Molecular convergence between lineages with low susceptibility to ADB

Using a pipeline for the detection of molecular convergence (grand-conv), we analysed 4,300 orthologous groups (OGs; supplementary table S1) that contained sequences from the nuclear genomes of five diploid ADB-susceptible *Fraxinus* species, three outgroup species, and at least two out of three lineages with low susceptibility to ADB (*F. mandshurica*, *F. platypoda* and three closely related species in section *Ornus* - *F. ornus*, *F. paxiana* and *F. sieboldiana*). Of these OGs, 2,904 included sequences from all 13 species. We conducted three pairwise comparisons between the lineages with low susceptibility, running grand-conv on multiple sequence alignments from the set of OGs encompassing each pair of lineages in turn (see Materials and Methods). This identified 68 loci containing at least one amino acid variant with a posterior probability of convergence of ≥0.90 and where the evidence of excess convergence (i.e. in excess of that expected due to chance alone given the level of divergence observed between lineages) was highest between lineages with low susceptibility to ADB. After filtering for false positives resulting from paralogy (including where this could not be discounted with certainty - see Materials and Methods), gene model errors and alignment errors, 62 loci containing 69 amino acid sites with evidence of convergence were retained (supplementary table S2).

Signatures resembling molecular convergence might also arise from introgressive hybridization or incomplete lineage sorting (ILS). To check for this possibility, we examined gene-trees for the hierarchical orthologous group (HOG) of each locus, which included sequences from all taxa (encompassing 29 *Fraxinus* genome assemblies and three outgroups - see Materials and Methods and Figure 1a). Discounting loci showing evidence for the inclusion of paralogous sequences within the OG (see above), we found only three cases where the sequences with convergent amino acid residues grouped more closely within the HOG gene-trees than expected from the species-tree (supplementary Figure 1a, c, e, albeit without strong support). For these loci, we repeated gene-tree inference following the exclusion of sites where convergence had been detected (supplementary table S2) and found the sequences that had contained convergent character states no longer clustered in a single group, with the topology more closely reflecting the species-tree (supplementary Figure 1b, d, f). Therefore, for all 62 loci within our filtered set, the pattern of shared amino acid states between lineages with low susceptibility to ADB was better explained by convergent mutations than the alternative hypotheses of introgressive hybridization or ILS.

**Fig. 1.**
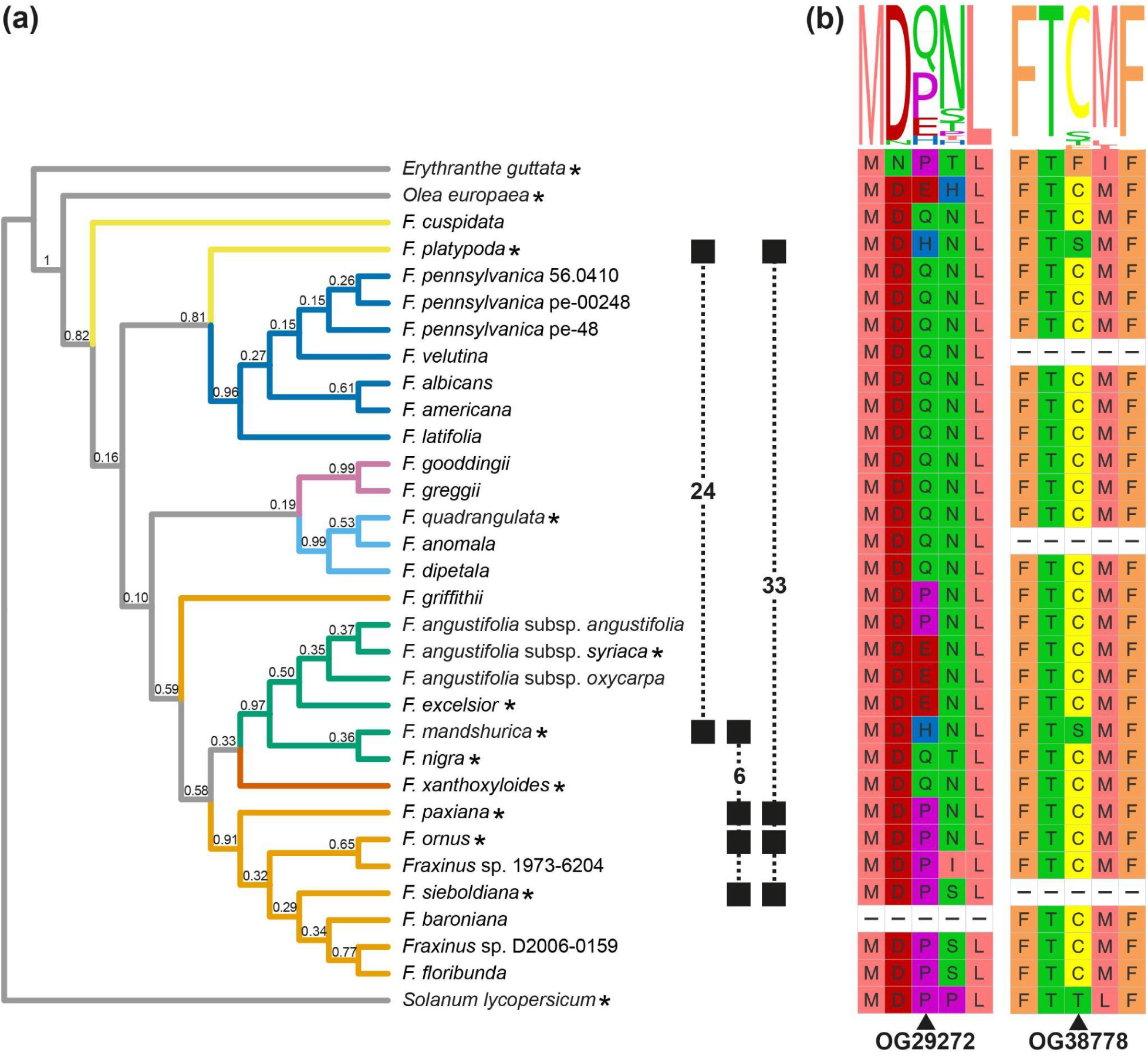
Evidence of molecular convergence between lineages with low susceptibility to ADB. a) Number of candidate loci detected for each of the three pairwise comparisons, shown in the context of a species-tree for *Fraxinus* inferred from 267 phylogenetically informative loci found in all taxa via Bayesian concordance analysis. Taxonomic sections within *Fraxinus* according to Wallander (2012) are shown in different colours; light blue, section *Dipetalae*; green, section *Fraxinus*; dark blue, section *Melioides*; light orange, section *Ornus*; pink, section *Pauciflorae*; dark orange, section *Sciadanthus*. *Fraxinus* species not placed in a specific section (*incertae sedis*) are coloured yellow, and outgroups grey. Numbers above the branches are sample-wide concordance factors. Asterisks following species names indicate the set of 13 taxa included in the convergence analyses. Filled squares (linked by dashed lines) denote taxa with low susceptibility included in the three pairwise analyses of molecular convergence, with the number of candidates found from that comparison shown; numbers do not sum to 62 (that is, the total number of candidate genes) because one locus was identified by two comparisons. b) Example loci showing patterns of strict convergence between lineages with low susceptibility to ADB, both identified from the *F. mandshurica* versus *F. platypoda* pairwise comparison. Sites at which evidence of convergence was detected (indicated with arrowheads) are shown with the two flanking amino acid positions on either side, based on alignments of full-length protein sequences including all taxa: left - OG29272 positions 1-5; right - OG38778 positions 88-92. Sequence rows are presented according to the order of taxa in the species-tree in part a); dashes indicate taxa which lack gene model predictions for these loci. Sequence logos indicating the degree of conversation across taxa are shown above the multiple sequence alignment segments (visualisations generated using the ggmsa package (Zhou et al. 2022) v 1.8.0 in R; amino acids coloured according to their physicochemical properties using the Zappo_AA scheme). Alt text: Graphical summary of evidence for amino acid convergence between *Fraxinus* species with low susceptibility to ash dieback disease, showing a phylogenetic tree with overall numbers of candidate loci detected between three pairs of *Fraxinus* lineages and example segments of multiple sequence alignments for loci with loci with strict convergence.

Fifty five of the 62 loci contained a single putative convergent site, with two sites detected in each of the remaining seven loci (supplementary table S2). The largest number of loci was detected from the comparison between *F. platypoda* and species from section *Ornus* (33 loci; Figure 1a), followed by the *F. mandshurica* versus *F. platypoda* comparison (23 loci), with the fewest candidate loci (five) detected from the *F. mandshurica* versus section *Ornus* comparison. A further locus was identified from two of three pairwise comparisons (Figure 1a; supplementary table S2) and thus can be considered particularly well supported. When considering the full set of 29 *Fraxinus* genome assemblies (which includes taxa where the level of susceptibility to ADB is unknown or uncertain, as well as further highly susceptible taxa), and not just those used for convergence analyses, for the majority of loci the amino acid states inferred as convergent between species with low susceptibility to ADB were also found in additional taxa (supplementary table S2). However, for eight loci there was a pattern of strict convergence, where the convergent amino acid state is only present in the lineages with known low susceptibility: OG4870, OG8392, OG8679, OG9544, OG14715, OG29272, OG38778 and OG44379 (supplementary table S2, Figure 1b). We note, however, that our sample size for the majority of species was a single genome assembly, that is not haplotype resolved, and we cannot exclude the possibility that candidate alleles are segregating undetected within species.

### Putative functions of candidate loci

We performed functional annotation and pathway analysis for the *F. excelsior* reference gene models for our 62 candidate loci and inferred their orthologues in *Arabidopsis thaliana* and *Populus trichocarpa*. Fifty six of the 62 candidate genes had at least one GO term annotated, with 45 annotated with at least one pathway (supplementary table S2). Probable *A. thaliana* and *P. trichocarpa* orthologues were identified for 43 and 48 of the candidate loci, respectively. Of 19 loci lacking evidence for an *A. thaliana* orthologue, close paralogues (i.e. genes belonging to the same HOG as the candidate locus) could be inferred for nine, and for 14 loci lacking a *P. trichocarpa* orthologue, close paralogues were identified for eight (supplementary table S2). Based on the GO term and pathway annotations, and evidence for the function of orthologues or close paralogues in *A. thaliana* and *P. trichocarpa*, 20 (32%) of the *Fraxinus* candidate loci can be linked to immunity or defence response (table 1), with a further three loci (5%) having a possible link to these functions (supplementary table S2). Of the 20 loci linked to immunity or defence response, 11 have evidence for a role in defence against fungal pathogens specifically (table 1).

**Table 1.** Candidate loci linked to immunity or defence response.

| Candidate locus | *Fraxinus excelsior* gene model | Convergent change(s) detected | *Arabidopsis thaliana* orthologue or close paralogue(s) | Selected GO terms for *F. excelsior* and/or *A. thaliana* genes | Pathway annotation for *F. excelsior* gene | External evidence for fungal defence response related function? |
| --- | --- | --- | --- | --- | --- | --- |
| OG2715 | FRAEX38873_v2_000309880 | A17V; T920S | AT1G64060 | FRAEX38873_v2_000309880: defense response by callose deposition, respiratory burst involved in defense response.<br>AT1G64060: regulation of defense response, defense response to fungus, defense response to other organism. | MAPK signaling pathway - plant; plant-pathogen interaction; generation of superoxide radicals. | Yes |
| OG4171 | FRAEX38873_v2_00<br>0334500 | N97K; L270V | AT4G21390 | AT4G21390: immune system process, regulation of defense response, defense response to bacterium, defense response to fungus. | MAPK signaling pathway; NF-kappa B signaling pathway; NOD-like receptor signaling pathway; Toll-like receptor signaling pathway; neurotrophin signaling pathway; toll and Imd signaling pathway; recognition of fungal and bacterial pathogens and immunity response. | Yes |
| OG4870 | FRAEX38873_v2_00<br>0276050 | K539R | AT2G24230 | FRAEX38873_v2_000276050: defense response to fungus. | Anther and pollen development | Yes |
| OG12319 | FRAEX38873_v2_00<br>0213420 | K358E | AT1G60420 | n/a | n/a | Yes |
| OG14715 | FRAEX38873_v2_00<br>0391250 | T311M | n/a | FRAEX38873_v2_000391250: defense response to bacterium, innate immune response. | n/a | No |
| OG14769 | FRAEX38873_v2_00<br>0396280 | N185K | AT3G15200 | AT3G15200: regulation of<br>defense response, defense<br>response to other organism. | Cytosolic glycolysis; lysine<br>biosynthesis II; lysine<br>biosynthesis I. | No |
| OG17061 | FRAEX38873_v2_00<br>0049880 | I349M | AT5G16820* | n/a | Longevity regulating<br>pathway - worm; HSFA7/<br>HSFA6B-regulatory<br>network-induced by drought<br>and ABA. | No |
| OG27529 | FRAEX38873_v2_00<br>0177770 | E19G | AT2G41880 | AT2G41880: immune<br>system process, regulation<br>of defense response,<br>defense response to<br>bacterium, defense<br>response to fungus. | Purine metabolism | Yes |
| OG28023 | FRAEX38873_v2_00<br>0270940 | I194V | AT3G46620 | AT3G46620: immune<br>system process, regulation<br>of defense response,<br>defense response to bacterium, defense response to fungus. | Methylerythritol phosphate<br>pathway | Yes |
| OG29272 | FRAEX38873_v2_00<br>0147540 | Q3H | AT5G42440 | AT5G42440: regulation of immune system process, regulation of defense response, defense response to bacterium, defense response to fungus. | MAPK signaling pathway; NF-kappa B signaling pathway; NOD-like receptor signaling pathway; Toll-like receptor signaling pathway; Neurotrophin signaling pathway; Toll and Imd signaling pathway; anther and pollen development. | Yes |
| OG29496 | FRAEX38873_v2_00<br>0009040 | A11D; G225A | AT1G75500* | n/a | Intracellular auxin transport | Yes |
| OG30871 | FRAEX38873_v2_00<br>0214730 | L184S;<br>S190P | AT1G70170 | AT1G70170: defense response to bacterium. | Recognition of fungal and bacterial pathogens and immunity response | Yes |
| OG31015 | FRAEX38873_v2_00<br>0183110 | H269D | AT3G61440 | AT3G61440: immune response. | Sulfur metabolism; cyanoamino acid metabolism; cysteine and methionine metabolism;<br>cysteine biosynthesis I. | Yes |
| OG33840 | FRAEX38873_v2_00<br>0384380 | L248S | AT4G23730 | AT4G23730: defense<br>response to other organism. | Glycolysis /<br>gluconeogenesis. | No |
| OG35529 | FRAEX38873_v2_00<br>0282600 | W137G | AT1G03610 | AT1G03610: defense<br>response to bacterium. | n/a | No |
| OG38778 | FRAEX38873_v2_00<br>0166900 | C90S | AT2G37940 | AT2G37940: immune<br>system process, defense<br>response to bacterium. | Sphingolipid signaling<br>pathway; sphingolipid<br>metabolism | Yes |
| OG45768 | FRAEX38873_v2_00<br>0158130 | H149N | AT4G05060*,<br>AT4G21450*,<br>AT5G54110* | AT5G54110: defense<br>response to other organism. | Flavin biosynthesis | No |
| OG50989 | FRAEX38873_v2_00<br>0237270 | S19P | AT4G26470 | n/a | Plant-pathogen interaction | No |
| OG51604 | FRAEX38873_v2_00<br>0300650 | G207S | AT1G32690 | AT1G32690: defense<br>response to other organism. | n/a | Possibly |
| OG55857 | FRAEX38873_v2_00<br>0174030 | S90Y | AT5G38700 | AT5G38700: regulation of<br>defense response, defense response to other organism. | n/a | Possibly |

In addition, we tested for overlap between our 62 candidate loci and a set of differentially expressed genes (DEGs) from a study of transcriptional response to *H. fraxineus* in *F. excelsior (Sahraei et al. 2020)*; these DEGs were detected from a comparison between tissues that were symptomatic and proximal to a necrosis or non-symptomatic and distal to the necrosis in plant materials infected with *H. fraxineus* and indicate genes that may function in response to ADB infection (Sahraei et al. 2020). We could assign a gene model from the *F. excelsior* reference annotation (BATG0.5) to 897 of the 1,009 DEGs (88.9%; see Materials and Methods and supplementary table S3). Two of our candidates (OG26849 and OG27529) are reciprocal best BLAST hits for a DEG, suggesting they may be involved in the transcriptional response to the fungus (supplementary tables S2 and S3), with one of these also among the 11 loci linked to fungal defence response on the basis of other evidence (table 1).

As well as roles directly involving immunity and defence response, evidence for other functions relevant to ADB were detected among our candidates. This includes two loci with potential roles in leaf senescence (OG30871 and OG38851), the timing of which has been found to be negatively correlated with the level of damage from ADB (McKinney et al. 2011), and six loci (OG2715, OG12857, OG26849, OG29496, OG35954 and OG48667) suggested to be involved in functions related to the cell wall (such as cell wall organisation and biogenesis; supplementary table S2), the penetration of which is part of the early infection process of *H. fraxineus* (Cleary et al. 2013). Three of these eight genes are also linked to immunity or defence response and one overlaps with those matching DEGs (see above). Together, genes with links to immunity or defence response (including those with possible links), leaf senescence or functions related to the cell wall account for 28 (45%) of our 62 candidates.

### Comparison with previously detected candidate loci from F. excelsior

Previous studies have reported loci associated with differing susceptibility to ADB in *F. excelsior* (Sollars et al. 2017; Stocks et al. 2019; Sahraei et al. 2020). Of 1,125 separate genes differentially expressed between *F. excelsior* genotypes with low and high ADB susceptibility in Sahraei et al. (2020), 869 (77.2%) could be matched to a BATG0.5 gene model (supplementary table S4). Of these 869 genes, two overlap with our candidate loci (OG2715 and OG4171; Figure 2 and supplementary tables S2 and S4). Both DEGs were detected from analysis of asymptomatic tissue of individuals with varying susceptibility to ADB that were inoculated with *H. fraxineus* (we could assign a BATG0.5 gene model to 840 of the 1,082 DEGs from this analysis, and to 84 of the 138 DEGs from an equivalent analysis of symptomatic tissue (Sahraei et al. 2020); Figure 2, supplementary table S4); these represent different DEGs to those found from a comparison of symptomatic and non-symptomatic tissues that overlap with two of our loci (see *Putative functions of candidate loci* above). In addition, one of the gene expression markers (GEMs) from Sollars et al.(2017) matches a candidate from our analyses (OG38851), whilst none of the 55 genes identified from the GWAS carried out by Stocks et al. (2019) overlap with our final set of loci (Figure 2). All three candidate loci that overlap with those from previous studies are included among the 28 loci with evidence for functions that are relevant to ADB. In addition to studies considering candidate genes, Nemesio-Gorriz et al (2020) identified 64 metabolites associated with level of susceptibility to ADB in *F. excelsior*, belonging to nine biochemical classes and which they propose contribute to constitutive chemical defence against *H. fraxineus* (see Materials and Methods). None of our candidate loci are suggested to play a role in the biosynthesis, catabolism or metabolism of these metabolites.

**Fig. 2.**
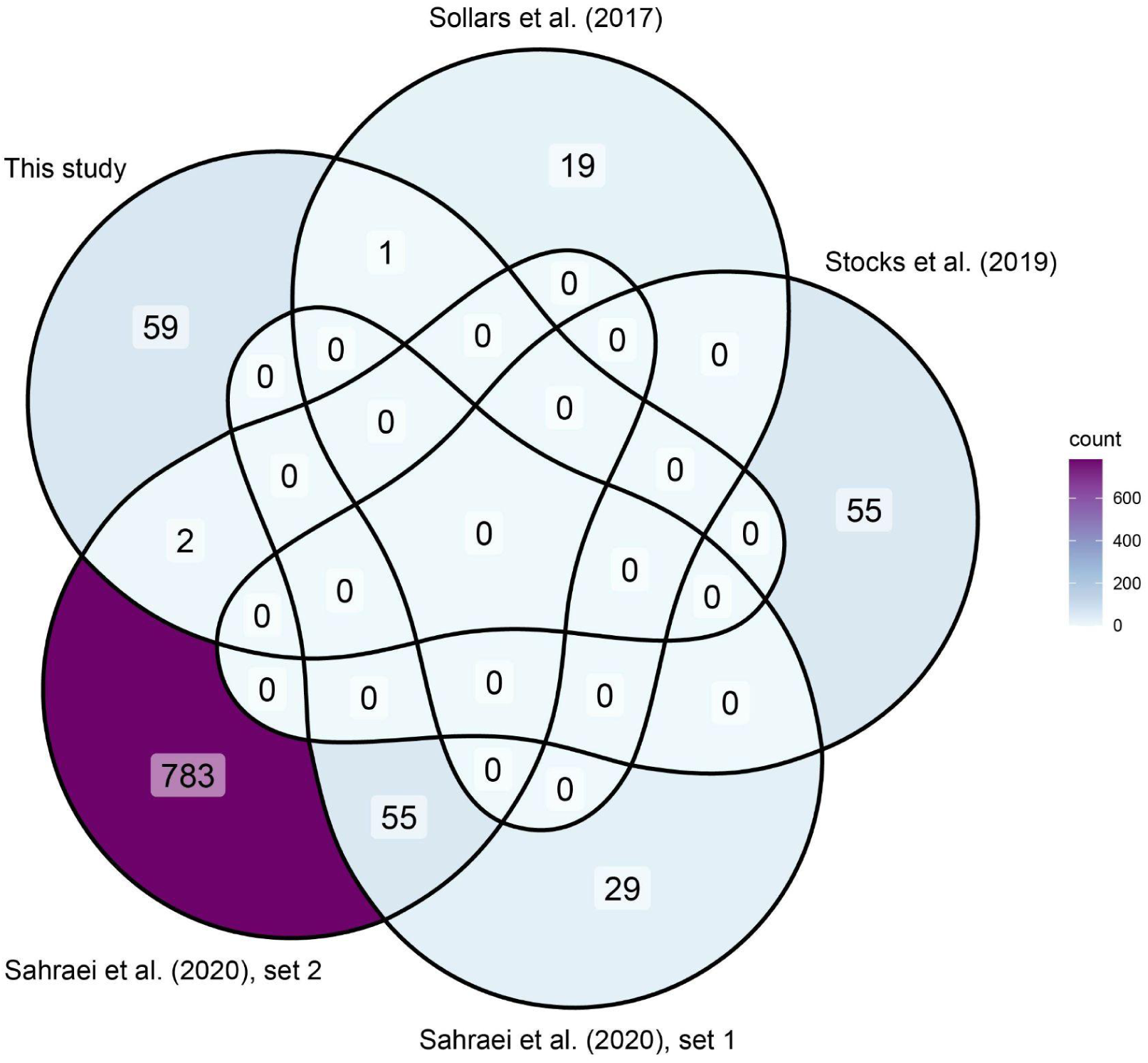
Venn diagram showing overlap between candidate loci identified via a molecular convergence approach and those associated with level of susceptibility to *H. fraxineus* in *F. excelsior* identified from previous analyses. Gene sets identified in previous studies are: Sollars et al. (2017) - 20 gene expression markers (GEMs) identified via associative transcriptomics; Stocks et al. (2019) - 55 genes containing or within 5 kb up or downstream of a SNP significantly associated with health status in a GWAS; Sahraei et al.(2020), set 1 - 84 genes with evidence of differential expression between inoculated *F. excelsior* individuals with varying levels of susceptibility, identified from sequencing of symptomatic tissue; Sahraei et al.(2020), set 2 - 840 genes with evidence of differential expression between inoculated *F. excelsior* individuals with varying levels of susceptibility, identified from sequencing of asymptomatic tissue. See main text section *‘Comparison with previously identified candidate loci’* for further details of the gene sets from these studies. Alt text: Venn diagram summarising numbers of candidate loci for low susceptibility to ash dieback disease shared between the current study and studies published previously.

## Discussion

Using genomic data from multiple *Fraxinus* species with both high and low susceptibility to the ADB causing fungus, *H. fraxineus*, we have uncovered 62 loci containing 69 amino acid sites with evidence of convergence between known and putative natural hosts. We hypothesise these genes contribute to defence against this otherwise deadly pathogen in ash tree species that likely share a coevolutionary history with *H. fraxineus*, whilst acknowledging alternative possible explanations below.

### External evidence links candidate loci to defence response related functions

Eleven of the candidate loci detected from our molecular convergence analyses (see Table 1) have external supporting evidence to suggest they function in defence response to fungal pathogens. OG2715 (FRAEX38873_v2_000309880) is orthologous to AT1G64060, which is involved in production of reactive oxygen species within the plant immune response to pathogens (Morales et al. 2016). AT1G64060 expression is dependent on TGA transcription factors - central regulators of immunity genes (Qi et al. 2022). OG4171 (FRAEX38873_v2_000334500) is orthologous to AT4G21390, which encodes a G-type lectin - receptor kinases whose functions include defence response against pathogens (Teixeira et al. 2018). AT4G21390 is upregulated in response to several fungal pathogens (Chae et al. 2009) and differentially expressed under a range of abiotic stressors (Mondal et al. 2021). OG4870 (FRAEX38873_v2_000276050) is orthologous to AT2G24230, which encodes a leucine-rich repeat protein kinase family protein involved in protein phosphorylation (Nemoto et al. 2011). Our functional annotation of FRAEX38873_v2_000276050 indicates this gene is also specifically involved in ‘defense response to fungus’. OG12319 (FRAEX38873_v2_000213420) is orthologous to AT1G60420, which plays a role in protecting plant cells from oxidative stress resulting from the immune response following pathogen attack (Kneeshaw et al. 2017).

AT1G60420 is also involved in activating defence response to the fungal pathogen *Alternaria brassicicola* (Kang et al. 2020). OG27529 (FRAEX38873_v2_000177770) is orthologous to AT2G41880, which is involved in nucleotide metabolism (Kumar et al. 2000) and was annotated with multiple GO terms indicating a role in immunity, including ‘defense response to fungus’, ‘defense response to bacterium’ and ‘regulation of defense response’ (Depuydt & Vandepoele 2021). Moreover, FRAEX38873_v2_000177770 is differentially expressed in response to *H. fraxineus* infection in *F. excelsior* (Sahraei et al. 2020). OG28023 (FRAEX38873_v2_000270940) is orthologous to AT3G46620, which encodes a RING-type E3 ubiquitin ligase and positively regulates pattern triggered immunity and salicylic acid mediated defence signalling (Yi et al. 2022). OG29272 (FRAEX38873_v2_000147540) is orthologous to AT5G42440, a leucine rich repeat receptor-like kinase gene, (Diévart & Clark 2003) annotated with many terms suggesting a role in defence response, including ‘defense response to fungus’ (Depuydt & Vandepoele 2021). OG29496 (FRAEX38873_v2_000009040) appears to be a close paralogue of AT1G75500 (*WAT1*), which is required for secondary cell wall formation in *A. thaliana* but also plays a role in mediating resistance to bacterial and fungal vascular pathogens (Denancé et al. 2013). AT1G70170 is the orthologue of OG30871 (FRAEX38873_v2_000214730), and is a matrix metalloproteinase gene suggested to contribute to plant immune response. AT1G70170 expression is induced in response to infection by the fungal pathogen *Botrytis cinerea*, as well as to a bacterial pathogen (Zhao et al. 2017), and mutants overexpressing this gene have enhanced *B. cinerea* resistance (Zhao et al. 2013). It is noteworthy that this candidate is also inferred to play a role in the regulation of leaf senescence (supplementary table S2), with *A. thaliana* individuals carrying a knock-out mutation of AT1G70170 demonstrating earlier leaf senescence than wild type plants (Golldack et al. 2002). Earlier leaf senescence is observed to correlate with lower levels of damage from ADB in *F. excelsior (McKinney et al. 2011; Doonan et al. 2025)*. A proposed explanation for this relationship is that early leaf senescence may limit stem infection establishment, by reducing time for the fungus to grow from leaves into the woody tissues in a given growing season (Landolt et al. 2016), although *F. excelsior* shows evidence for an active defence response (McKinney et al. 2012). It has also been suggested the correlation could result from linkage between genes underlying these traits (Landolt et al. 2016). However, our results indicate an individual gene may contribute to the control of both. Another candidate (OG38851) is also linked to leaf senescence (supplementary table S2); the observation that numerous genes are involved in both plant immunity and leaf senescence has led to the suggestion these processes may have partially overlapping regulatory networks (Zhang et al. 2020). OG31015 (FRAEX38873_v2_000183110) is orthologous to AT3G61440, which encodes a cyanide detoxifying enzyme implicated in immune response, with loss of function mutants having increased susceptibility to a necrotrophic fungus and decreased susceptibility to a hemibiotrophic bacterium (García et al. 2013). Moreover, transcription of its orthologue in poplar (Potri.002G160800.1; supplementary table S2) was induced in response to infection by a hemibiotrophic fungal pathogen (Liao et al. 2021). Finally, OG38778 (FRAEX38873_v2_000166900) is orthologous to AT2G37940 which has been implicated in the regulation of programmed cell death associated with defence response to powdery mildew (Wang et al. 2008).

As well as the eleven genes explicitly linked to defence response to fungi, other notable loci include OG26849 (FRAEX38873_v2_000344290) and OG50989 (FRAEX38873_v2_000237270). OG26849 not only appears to play a role in cell wall organisation based on its orthologue in *A. thaliana* and results of our functional annotation (supplementary table S2), with relevance to both the mechanism of ADB infection (Cleary et al. 2013) and function of putative host interaction genes in *H. fraxineus* (McMullan et al. 2018), but also shows evidence for differential expression in response to *H. fraxineus* infection. OG50989 is suggested to be involved in the ‘plant-pathogen interaction’ pathway on the basis of our functional analyses (table 1) and is also a candidate gene for defence against the emerald ash borer (EAB; (Kelly et al. 2020). As some *Fraxinus* species with low susceptibility to ADB also share the trait of resistance to EAB (Kelly et al. 2020), our analyses targeted at identifying genes related to ADB may have simultaneously detected those relevant to EAB resistance, albeit the species with the highest susceptibility differ between these threats (Kelly et al. 2020). Alternatively, OG50989 may indeed contribute to defence against both ADB and EAB, given its likely function in calcium ion binding (supplementary table S2) and the fact calcium signalling plays a key role in plant immune responses to multiple organisms (Aldon et al. 2018).

Although many of our candidate genes have evidence for functions relevant to the response against ADB, more than half (34 out of 62; see Results) lack such evidence. These may include genes with roles in immunity or defence that are not yet understood (Depuydt & Vandepoele 2021), or that relate to other phenotypic traits correlated with low ADB susceptibility (see above). Whilst we took steps to control for false positives, we cannot entirely discount that some remain, although the analytical approach used is considered less prone to these than some alternatives (Macdonald et al. 2025). As outlined above (see Results) most amino acid states inferred as convergent between species with low susceptibility to ADB were found in additional taxa. Some of these might not represent convergent evolution, but rather segregation of ancestral standing genetic variation; inclusion of more taxa of known phenotype in future analyses could change the reconstruction of ancestral states and may be expected to reduce the number of candidate loci with evidence of repeated substitutions (Thomas et al. 2017). Increased intraspecific sampling would also enhance further tests of the strength of association between the detected amino acid variants and susceptibility to ADB. Even if ancestral variation, rather than convergent mutations, is the source of some variants we detect, it does not necessarily mean this has not been subject to natural selection in separate lineages (e.g. (Louis et al. 2021)) nor that it does not contribute to low ADB susceptibility.

It should be noted that not all instances of molecular convergence reflect adaptive evolution; some cases of identical amino acid changes may not imply a shared functional change since their impact can depend on the wider genetic background, although the likelihood of them having the same phenotypic effect is increased in closely related species (Storz 2016). We also acknowledge that potential functions of our candidate genes are mainly informed by evidence from *A. thaliana*, or other model species (see above and Table S2). The complexity of plant genomes, reflecting a history of whole genome multiplications, limits the usefulness of orthology for transferring gene functions from *A. thaliana* to other species since duplicated genes may diverge in function (Roeder et al. 2025) and trees and herbaceous plants differ in copy numbers for some disease resistance gene families (Birkeland et al. 2026). Thus, we cannot rule out that even candidates with strong evidence for roles in defence against fungal pathogens are actually performing other functions in *Fraxinus*. Whilst five of our candidates have evidence for expression in response to *H. fraxineus* specifically (OG2715, OG4171, OG26849, OG27529 and OG38851; see Results), future experimental validation of roles in defence against ADB for additional loci would be advantageous.

### Analysis of natural hosts detects novel candidate loci

Previous attempts to discover candidate loci for low susceptibility to ADB have mainly focused on variation within *F. excelsior*, a recently acquired host of *H. fraxineus* within Europe, either in terms of single nucleotide polymorphisms (SNPs; (Stocks et al. 2019; Doonan et al. 2025)) or gene expression differences between genotypes with varying susceptibility (Harper et al. 2016; Sollars et al. 2017; Sahraei et al. 2020). Stocks et al. (2019) detected 55 candidate genes that either contained a SNP significantly associated with health status of ADB-infected trees or which were within 5 kb either up or downstream of such a SNP. We had limited power to detect convergence in these genes since only eight were included in our analyses (supplementary Figure S2), none of which contained a significant SNP that alters the predicted protein sequence in *F. excelsior* (Stocks et al. 2019). Given this, the lack of overlap with our final set of candidates is unsurprising (Figure 2). Only four of these eight genes were detected as candidates with a new GWAS that reanalysed data from Stocks et al. (2019) using a pangenome approach (Wood et al. 2025), showing that even similar methodological approaches may yield differing sets of loci. We found greater overlap between our final set of candidates and loci detected from analyses of transcriptome data, with one GEM (Sollars et al. 2017) and two DEGs (Sahraei et al. 2020) common to our candidates (Figure 2). All three loci (OG2715, OG4171 and OG38851) also have external evidence for roles relevant to ADB (see above).

Despite finding some common genes, the minimal overlap between our candidates and those from *F. excelsior* suggests this naïve host may lack many of the variants that confer low susceptibility in species likely to have coevolved with *H. fraxineus*. Differences in the loci detected may be partially methodological, as noted above. We could not consider all *F. excelsior* genes in our analyses (see Fig. S2) - however, the genome-wide nature of previous analyses, and the fact that GWAS include genes that are thousands of base-pairs away from significant SNPs as candidates, means that if the loci we identify are associated with low susceptibility in *F. excelsior* we might expect those studies to have been able to detect them. It is notable that 211 candidate genes associated with low susceptibility to ADB from a recent *F. excelsior* pangenome-based GWAS also did not overlap with any of our loci (Wood et al. 2025), further indicating that there may be true inter-specific differences in the genomic basis of this trait. Given that *F. excelsior* is not believed to have shared a coevolutionary history with *H. fraxineus* prior to the fungus arriving in Europe within the past few decades, variants contributing to reduced susceptibility in this species are inferred to reflect the evolutionary response to other selection pressures (i.e. exaptation (Gould & Vrba 1982)), possibly including related species of fungi. Indeed, it has been suggested that the low susceptibility to ADB observed in some *F. excelsior* individuals may result from coevolution with *H. albidus* (Landolt et al. 2016), a native

European relative of *H. fraxineus*, and pleiotropic effects could lead to the same genes contributing to reduced susceptibility to multiple pathogens (Capador-Barreto et al. 2021). Another possibility is loci contributing to low susceptibility in *F. excelsior* could represent ancestral variation retained from a progenitor that was exposed to selection from *H. fraxineus* (or its progenitor), potentially the common ancestor with *F. mandshurica* (one of the extant natural hosts and a close relative of *F. excelsior* (Kelly et al. 2020; Hinsinger et al. 2013; Wallander 2012)). Similar explanations have been proposed to account for variation in susceptibility in apparently naïve hosts of other plant pathogens (Tobias et al. 2016). Such a scenario might explain the partial overlap between loci detected from our analyses, and those from studies on *F. excelsior*. Nevertheless, the majority of our candidate genes have not been detected from previous analyses, emphasising the complementary nature of studies focusing on coevolved versus naïve hosts.

### Implications for efforts to restore tree populations

Resistance breeding programmes have been established for a number of ecologically and economically important tree species that are threatened by non-native pests and pathogens (Sniezko & Liu 2021). Such programmes can provide plant materials to facilitate restoration of damaged forest ecosystems (Sniezko & Koch 2017). The majority of resistance breeding programmes have attempted to leverage variation in disease susceptibility found within the affected tree populations, which typically represent newly acquired hosts within a recently invaded region (Sniezko & Koch 2017). Hybridisation between related species with differing susceptibility has also been used to introduce beneficial alleles into species under threat, for example with breeding for reduced susceptibility to chestnut blight and Dutch elm disease (Woodcock et al. 2019). Knowledge of the genomic basis of resistance or susceptibility in natural hosts that have coevolved with a pest or pathogen may be useful for the development of tools for marker assisted selection, to guide the introgression of alleles into the target species (Sniezko et al. 2023).

In the case of the ADB epidemic in Europe, efforts to date have focused on breeding within *F. excelsior* (Langer et al. 2022; Stocks et al. 2019). The new candidate loci we identified in other ash species could help inform these efforts, including via targeted analysis of these genes to explore variation within *F. excelsior* (for example, to seek rare beneficial alleles), or by guiding strategies for genome editing. The partial overlap between loci detected from Asian species of ash and those previously identified within *F. excelsior* may also increase confidence in the latter. Our loci could also be used to facilitate breeding programmes involving hybridisation. Several wide hybrids between different taxonomic sections of *Fraxinus* have been reported, including between members of section *Ornus* (which contains multiple Asian species with low susceptibility to ADB) and *F. excelsior* ((Plumb et al. 2025) and references therein). This suggests the transfer of alleles from *Fraxinus* species with low susceptibility into those that are highly susceptible may be achievable.

The nature of our analyses, by definition, limited us to considering amino acid variation shared across multiple lineages. Thus, much of the genetic basis of low susceptibility to ADB in the species we studied may remain to be discovered, whether it be due to lineage-specific alleles, structural variants, variants in regulatory regions, or other types of genomic change (Allard & Kumar 2026). Moreover, the binary classification of taxa as having either low or high susceptibility represents a simplification of the actual variation observed (e.g. see (Nielsen et al. 2017)) and may have caused certain patterns of amino acid convergence to be missed by our analyses. A more detailed and precise characterisation of susceptibility to ADB across the genus (such as through replicated common garden experiments) could increase statistical power to detect associated genetic changes, especially if multiple and varying resistance mechanisms are involved (Macdonald et al. 2025) as has been posited for *F. excelsior* (Schertler et al. 2026). This study therefore likely represents the tip of the iceberg of genomic resources present in the genus *Fraxinus* that could be deployed to help combat ADB, and emphasizes the wealth of relevant genetic variation that remains untapped within hosts from the pathogen’s native range. A similar approach to that used here could be applied to other major tree health threats to identify candidate genes to facilitate restoration efforts, especially in cases where their coevolved hosts span multiple independent lineages.

## Materials and Methods

### Taxon selection

For our analyses of evidence for molecular convergence, we made use of an existing genomic dataset for 22 diploid *Fraxinus* species (represented by 29 genome assemblies), spanning all major infrageneric taxonomic groups and almost half of the 48 species recognised by Wallander (2012); see Kelly et al. (2020) for further details. From among the taxa included in this dataset, we identified a subset representing those with both low and high susceptibility to ADB for inclusion in the convergence tests themselves, based on levels of damage/severity of symptoms reported for these taxa in the literature. Symptoms of ADB damage commonly used to estimate susceptibility include crown dieback, presence of epicormic shoots and necroses/lesions on stems and leaves (Kirisits & Schwanda 2015; Nielsen et al. 2011; Drenkhan, Solheim, et al. 2017; Drenkhan, Agan, et al. 2017; McKinney et al. 2014). There is no universally adopted standardised protocol for assessment of ADB susceptibility; approaches used include visual assessment of overall damage in younger trees on a categorical scale (with the number of categories varying by study), visual assessment of crown health in mature trees (as a percentage dead/alive, with fixed categories/health classes sometimes applied) and the measurement of lesion lengths from artificial stem inoculations (e.g (Nielsen et al. 2011; Hiemstra et al. 2019; Gross & Sieber 2016; Drenkhan, Agan, et al. 2017)). We took a consensus view of published results to determine which species tend to show the lowest and highest levels of damage from ADB overall, since the variation in methods used does not allow a more precise determination of susceptibility. Consequently, although susceptibility to ADB is a quantitative trait, for the purposes of our analyses we treat it as binary, placing taxa with minimal reported symptoms when exposed to ADB (e.g. comparatively low crown dieback, no or small lesions relative to other species measured) in the “low susceptibility” group and those with severe symptoms (e.g. comparatively high crown dieback, large lesions relative to other species) in the “high susceptibility” group.

We selected five *Fraxinus* species reported as having a low level of susceptibility to ADB (Drenkhan, Agan, et al. 2017; Nielsen et al. 2017; Drenkhan, Solheim, et al. 2017; Cleary et al. 2016; Queloz et al. 2017; Hiemstra et al. 2019; Kirisits & Schwanda 2015), and which belong to three separate well-supported clades within the species-tree for the genus (Kelly et al. 2020): *F. mandshurica* (the only confirmed diploid natural host of *H. fraxineus*), *F. ornus*, *F. paxiana*, *F. platypoda* and *F. sieboldiana*. These species are native to East Asia and the Russian Far East (within or close to the non-invasive distribution of *H. fraxineus* (Marçais et al. 2022)), except *F. ornus*, which occurs in Europe and Western Asia (Wallander 2012) but belongs to a predominantly East Asian clade (Wallander 2012; Kelly et al. 2020).

We also selected five taxa reported to have higher levels of susceptibility (Drenkhan, Agan, et al. 2017; Nielsen et al. 2017; Gross & Sieber 2016; McKinney et al. 2014; Coker et al. 2019; Hiemstra et al. 2019): *F. angustifolia* subsp. *syriaca* (susceptibility sometimes reported under the name *F. sogdiana*, which Wallander (2012) treats as a synonym of *F. angustifolia* subsp. *syriaca*), *F. excelsior*, *F. nigra*, *F. quadrangulata* and *F. xanthoxyloides*. These taxa are native to Europe, Western Asia, Central Asia, North America and North Africa, with *F. xanthoxyloides* also extending into the Himalaya (Wallander 2012). We also note that, whilst there are rare *F. excelsior* individuals which are relatively unaffected by ADB (see Introduction), the genomic data included in our analyses are from a susceptible individual (Sollars et al. 2017).

Finally, we included three outgroups: *Olea europaea* (from the same family as *Fraxinus* (Oleaceae)), *Erythranthe guttata* (from the same order (Lamiales)) and *Solanum lycopersicum* (from the same major eudicot clade (lamiids)). By limiting convergence tests to this subset of taxa we maximised the number of genes included in our analyses, as with increased taxa the number of genes annotated in all species decreases due to the fragmented nature of the genome assemblies (Kelly et al. 2020).

### Analysis of convergent evolution

To detect loci with evidence of molecular convergence at the amino acid level, we used sets of putatively orthologous sequences (OGs) previously generated by Kelly et al. (2020) with OMA standalone v.2.0.0 (Altenhoff et al. 2019), encompassing data from the 29 *Fraxinus* genome assemblies and three outgroups. Three pairwise comparisons were used when screening for loci with evidence of amino acid convergence between taxa with low susceptibility to ADB:

1. *F. mandshurica* (section *Fraxinus*) versus *F. ornus*, *F. paxiana* and *F. sieboldiana* (section *Ornus*)
2. *F. mandshurica* versus *F. platypoda* (*incertae sedis*)
3. *F. platypoda* versus *F. ornus*, *F. paxiana* and *F. sieboldiana*

These were all analysed in the context of the five susceptible species and the three outgroups (see *Taxon selection*). OGs that included protein sequences for all 13 taxa chosen for inclusion in the convergence analyses, or for one of the subsets of 10–12 taxa for each of the three pairwise comparisons outlined above, were selected from those inferred by Kelly et al (2020). The number of OGs included in each comparison was 3,124 OGs in analysis 1, 4,203 OGs in analysis 2 and 3,021 OGs in analysis 3, representing 4,398 unique loci, with 2,975 included in all three analyses. Sequences corresponding to preliminary gene models annotated within the *F. excelsior* organellar genomes (FRAEX38873_v2_000400370 - FRAEX38873_v2_000401330, which were included in file

Fraxinus_excelsior_38873_TGAC_v2.longestCDStranscript.gff3 (available from: https://doi.org/10.5281/zenodo.6948952) used by Kelly et al. (2020) for genome annotation of the other ash species (2020), but not reported in the reference genome publication (Sollars et al. 2017)), were included in the OMA analysis. However, none of the OGs containing sequences for these gene models were among those analysed here and all genes that were analysed are considered to be nuclear. In addition, file Fraxinus_excelsior_38873_TGAC_v2.longestCDStranscript.gff3 includes 91 gene models annotated on reference genome contigs (Sollars et al. 2017) that were later determined to be from contaminating organisms (Stocks et al. 2019). Three of these gene models (FRAEX38873_v2_000008270, FRAEX38873_v2_000220000 and FRAEX38873_v2_000391870) were included in the convergence analyses, but as they belong to OGs that also contain sequences from all three outgroups, and the *Fraxinus* taxa of interest, they seem unlikely to be true contaminants; none were among our final set of candidate loci.

Multiple sequence alignments of the CDSs for the OGs were generated using GUIDANCE2 (Sela et al. 2015) as described in Kelly et al. (2020). For the 2,975 loci shared across the three pairwise comparisons, a single alignment containing all 13 taxa was generated, whilst alignments for loci unique to a given analysis (the remaining 1,423 loci, as none were shared between just two comparisons) included the taxa for that pairwise comparison only. Alignments were filtered as described in Kelly et al.(2020), with the exception that those of <300 characters in length or that included sequences with <50% non-gap characters were excluded from further analysis, using the filter_alignments_on_length_and_missing_data.pl script available from: https://github.com/lkelly3/eab-ms-scripts. Following filtering of the alignments, the number of OGs remaining for input into the convergence pipeline itself was: 3,052 OGs for analysis 1, 4,107 OGs for analysis 2 and 2,949 OGs for analysis 3, representing 4,300 unique loci, with 2,904 included in all three analyses (supplementary table S1).

Multiple sequence alignment files were formatted as described in Kelly et al.(2020) and Grand-Convergence v.0.8.0 (hereafter referred to as grand-conv) used to detect loci with significant evidence for molecular convergence, in excess of that expected due to chance alone given the level of divergence observed (https://github.com/dekoning-lab/grand-conv). For the 2,904 OGs included in all three pairwise comparisons, sequences from the taxa not required for a particular analysis were removed from the alignment files prior to running grand-conv, using Unix commands. The grand-conv pipeline was run as described in Kelly et al. (2020) with Newick format species-tree files for each of the pairwise comparisons created by pruning the primary concordance tree from Kelly et al. (2020) to retain only the taxa that were relevant to that analysis; trees were rooted on *S. lycopersicum*. Output files from grand-conv were filtered to retain loci with at least one amino acid site where the posterior probability of convergence exceeded ≥0.9000, using the filter_grand_convergence_results.pl script available at https://github.com/lkelly3/eab-ms-scripts. For loci passing this threshold, we calculated the convergence regression residuals from the indexData.js files generated by grand-conv using a custom Python script (available from: https://doi.org/10.5281/zenodo.20412204) and selected for further analysis OGs where the highest excess convergence was found for the low susceptibility branch pair.

This set of loci was further filtered to remove those where orthology/paralogy conflation, incomplete lineage sorting (ILS) or introgressive hybridisation could not be excluded as a possible cause of the signal of convergence detected by grand-conv. For each of the candidate loci, we identified the hierarchical orthologous group (HOG) from the OMA analysis to which they belonged and inferred their gene trees using MrBayes v.3.2.6 (Ronquist et al. 2012) as described in Kelly et al. (2020). Consensus gene-trees were checked for evidence that the sequences within which convergence was initially detected (i.e. those included in the grand-conv analyses) represent multiple paralogues, with sequences containing the ‘convergent’ state and those containing ‘non-convergent’ state belonging to separate clades. In addition, we checked for other patterns of incongruence with the species tree topology involving the sequences included in the convergence analysis that might indicate ILS or introgression. For several loci, the sequences carrying the putatively convergent states were clustered more closely than expected on the basis of the species-tree topology, suggesting that one of the processes outlined above could have confounded the convergence analysis. However, such patterns could also result from shared convergent amino acid states causing sequences to appear more closely related than is actually the case. Thus, for these loci, the gene-tree analysis was repeated with the amino acid sites at which convergence had been detected excluded. No loci had gene-tree topologies that indicated evidence for confounding processes following removal of the apparently convergent sites, but we excluded from further analysis one locus for which the MrBayes analysis failed to converge after 10 million generations (i.e. with an average standard deviation of split frequencies ≥ 0.01).

To identify the positions of sites with evidence of convergence within the full-length protein sequences, and to check for the presence of putatively convergent amino acid states in additional species, we generated multiple sequence alignments for the OGs including all available taxa (i.e. including data from up to 29 *Fraxinus* genome assemblies and three outgroups - see above) using GUIDANCE2 with the following parameter settings: --program GUIDANCE --msaProgram MUSCLE –seqType aa –bootstraps 100. Alignments including all codons (i.e. no filtering for column alignment confidence) were visualised using AliView v.1.26 (Larsson 2014) or Mesquite v.3.6 (Maddison & Maddison 2018) and segments containing the convergent sites located using the site-specific information from grand-conv; we excluded from further consideration any OGs where there was an indication that gene model or alignment errors had caused a false-positive signal of convergence (e.g. such as the presence of truncated protein sequences for a subset of the taxa).

### Species-tree inference

We conducted a new species-tree analysis subsequent to the identification of loci with evidence of convergent evolution, to account for the possible confounding effect of molecular convergence on the inference of the tree topology. Species-tree inference was carried out using full-length gene sequences with BUCKy v.1.4.4 (Ané et al. 2007; Larget et al. 2010), as described in Kelly et al. (2020), with the exception that the analysis was only run once, with the *a* parameter set to 1 (since previous tests found only minor differences between results with different *a* values (Kelly et al. 2020)) and, that as well as excluding three loci with evidence of molecular convergence between EAB-resistant taxa (OG2897, OG4469 and OG16673; Kelly et al. (2020)), we also excluded from the 272 phylogenetically informative loci on which the main species-tree analysis reported in Kelly et al. (2020) was based, two loci (OG13185 and OG14351) that were among our filtered set of candidates with evidence of convergence. The primary concordance tree obtained from this analysis was identical to that inferred from the set of 272 loci by Kelly et al. (2020), with only minor (i.e. 0.01) differences to the concordance factors (Figure 1a), therefore confirming that the topologies of the species-tree files used as input for grand-conv (see above - *Analysis of convergent evolution*) were appropriate. See supplementary file S1 for a list of 267 loci used.

### Functional annotation

To provide insight into putative functions of the candidate loci identified from the convergence analyses, and account for updated understanding of plant gene functions since the previous functional annotation (Sollars et al. 2017), we annotated the *F. excelsior* reference gene models using the functional analysis module in OmicsBox v.3.0.30 ((BioBam 2019). BLAST2GO (Götz et al. 2008), EggNOG-mapper (Huerta-Cepas et al. 2019) and InterProScan (Jones et al. 2014) were all used for annotation. Protein sequences from Sollars et al. (Sollars et al. 2017) (file Fraxinus_excelsior_38873_TGAC_v2.longestCDStranscript.gff3.pep.fa, available from: https://doi.org/10.5281/zenodo.6948952) were used as input, following removal of preliminary organellar gene models (see above - *Analysis of convergent evolution*).

For annotation with BLAST2GO, BLASTP was first run via the Diamond BLAST option (Buchfink et al. 2021), searching against the NR database (v.2023-02-01) in the ‘more sensitive’ sensitivity mode and with the following parameter settings: taxonomy filter set to only include Viridiplantae in the search; e-value threshold of 1e-5; ‘Optimize for Long Query Seqs’ off, number of BLAST hits set to 20, HSP length cutoff of 25 and no filter for ‘HSP-Hit Coverage’. GO mapping from the BLAST hits was performed with BLAST2GO with option ‘use latest version of database’, and GO annotation then run with the following settings: Filter GO by Taxonomy = Viridiplantae; E-Value-Hit-Filter = 1 (i.e. no further filtering on e-values, as all BLAST hits had a maximum e-value of 1e-5); HSP-Hit Coverage CutOff = 50; Hit Filter = 20 (i.e. no further filtering since the maximum number of BLAST hits per query was already 20); Only hits with GOs = off, and default settings for all other parameters. EggNOG-mapper v.2.1.0 with EggNOG v.5.0.2 was run with default settings. InterProScan was run using the default selection of member databases. The BLAST2GO and EggNOG-mapper annotations were merged using the ‘Merge EggNOG GO’ option from the GO Annotation menu, with a setting of 1e-5 for the ‘Seed Ortholog E-Value Filter’ and default setting for the ‘Seed Ortholog Bit-Score Filter’, the InterProScan annotations further merged with these using the ‘Merge GOs’ option within the InterProScan menu and ‘Validate GO Annotations’ from the GO Annotation menu run to remove any redundant GO terms.

We also used OmicsBox to run both a Gramene Pathway Analysis (using the Plant Reactome database) (Naithani et al. 2017) and KEGG Pathway Analysis (Kanehisa & Goto 2000) for the 62 candidate loci, using the protein sequences for the relevant *F. excelsior* gene models. The Gramene Pathway Analysis was run with the following settings: ‘Run BLAST to link via Protein IDs’ on; ‘Give Priority to Taxon’ off; BLAST e-value threshold of 1e-5 and default settings for all other parameters. The KEGG Pathway Analysis was run with the following settings: ‘Link KEGG Orthologs via EggNOG’ on; ‘Link via Enzyme Codes’ off; ‘Drug Development’ and ‘Human Diseases’ deselected under ‘Include Categories’.

We checked the final functional annotations of our loci to see if any indicated a role in the biosynthesis, catabolism or metabolism of nine classes of metabolites associated with level of susceptibility to ADB in *F. excelsior* (Nemesio-Gorriz et al. 2020): cinnamic acid esters, citric acid, coumarins, flavonoids, lignans, monolignols, phenylethanoids, secoiridoids (including iridoid glycosides) and shikimates. The following terms were searched for: cinnamic, citric, coumarin, flavonoid, iridoid, lignan, monolignol, phenylethanoid and shikimate.

### Further characterisation of candidate loci

To identify likely orthologues from *Arabidopsis thaliana* (arabidopsis) and *Populus trichocarpa* (poplar), we reran OMA standalone (using v.2.4.1 (Altenhoff et al. 2019)) on the set of protein sequences used for the previous OMA analysis conducted by Kelly et al. (2020) with the addition of datasets “Araport11_pep_20160703_representative_gene_model” (from arabidopsis.org) and “Ptrichocarpa_210_v3.0.protein_primaryTranscriptOnly.fa” (from phytozome-next.jgi.doe.gov) which contain protein sequences for a single transcript per gene for annotation versions Araport11 and v3.0 of the arabidopsis (Cheng et al. 2017) and poplar genome assemblies (Tuskan et al. 2006). We selected *A. thaliana* and *P. trichocarpa* for inclusion because they represent, respectively, the plant species and tree species with the most well characterised gene functions, and thus inference of orthologues from these species can provide more detailed insights into the putative functions of our candidate genes. For the species-tree (used during the inference of HOGs), we edited the final version of the species-tree used with OMA by Kelly et al. (2020) to manually add *A. thaliana* and *P. trichocarpa*. OMA was run with the AlignBatchSize parameter set to ‘10e6’, OutgroupSpecies set to ‘’Araport11_pep_20160703_representative_gene_model’, ‘Ptrichocarpa_210_v3’’, DoGroupFunctionPrediction set to “false”, the Newick format species-tree file specified via the SpeciesTree parameter, and default settings for all other parameters. Genes belonging to the same OMA group (OG) as a given *Fraxinus* gene were considered to be likely orthologues, whereas those absent from the OG but present in the same HOG as the *Fraxinus* candidate genes were considered to be close paralogues.

### Comparison with previously identified candidate loci

To determine whether our candidates match any found in previous studies of genome-wide or transcriptome-wide differences associated with the level of susceptibility to ADB in *F. excelsior*, we checked for overlap with gene models from different sets of loci, namely: genes containing or within 5 kb up or downstream of a SNP significantly associated with health status (Stocks et al. 2019); gene expression markers (GEMs) associated with ADB damage scores (Sollars et al. 2017); genes with evidence of differential expression between individuals inoculated with *H. fraxineus* and with varying levels of susceptibility, either from symptomatic and asymptomatic tissues (Sahraei et al. 2020). Two more recent GWAS for ADB susceptibility have also been published (Doonan et al. 2025; Meger et al. 2024), but neither detected loci that passed standard significance thresholds. Sollars et al. (2017) and Stocks et al. (2019) used the same *F. excelsior* assembly and gene annotation versions as used here (Sollars et al. 2017), allowing us to directly compare gene model names, although it should be noted the GEMs reported by Sollars et al. (2017) included four gene models excluded from the final gene annotation of BATG0.5 due to significant overlap and similarity with repetitive element sequences (FRAEX38873_v2_000048340, FRAEX38873_v2_000048360, FRAEX38873_v2_000054200 and FRAEX38873_v2_000251090) and which were therefore not considered in the other studies. Sahraei et al. (2020) based their analyses on an earlier version of the *F. excelsior* assembly (BATG0.4). Therefore, to find gene models from the BATG0.5 annotation (Sollars et al. 2017) that correspond to the differentially expressed genes (DEGs) from Sahraei et al. (2020), we conducted BLAST searches to identify reciprocal best hits. File Fraxinus_excelsior_38873_TGAC_v2.longestCDStranscript.gff3.cds.fa (available from: https://doi.org/10.5281/zenodo.6948952) was used for the BATG0.5 gene models and the fasta file provided in the supplementary information of Sahraei et al. (2020) for the DEGs. makeblastdb from BLAST+ v.2.2.31+ (Camacho et al. 2009) was used to format each file into a BLAST database and BLASTN searches performed in both directions with the following parameter settings: blastn -task blastn -evalue 1e-10 -max_target_seqs 1 -max_hsps 1 -outfmt 6. DEGs where a reciprocal best hit with a BATG0.5 gene model could be found were retained for further analysis, with the remainder of the DEGs (n = 514) discarded. Two of 91 gene models from contigs identified by Stocks et al. (2019) as likely deriving from contaminating organisms and therefore excluded from their analyses (see above - *Analysis of convergent evolution*) were among the retained set of DEGs from the analysis of asymptomatic tissue (FRAEX38873_v2_000328450 and FRAEX38873_v2_000391870), but no overlap was found with the other sets of loci. We retained these two genes in our comparisons because they form part of the final set of gene models reported in Sollars et al. (2017) and because there is some uncertainty over whether all of the 91 genes truly represent contaminant organisms (see above). Finally, three of the DEGs had a reciprocal best hit to one of the preliminary organellar gene models included in file Fraxinus_excelsior_38873_TGAC_v2.longestCDStranscript.gff3 (see above - *Analysis of convergent evolution*); we retained these in our comparisons because they were also considered in the other analyses.

The number of overlapping candidate loci between studies was summarised as a Venn diagram, generated using the ggVennDiagram package (Gao et al. 2024) (v.1.5.2) in R v.4.3.1 (R Core Team 2023). To summarise how many of the loci found by the previous studies we had the potential to detect, we also generated a Venn diagram showing overlap between the full set of gene model input into our convergence analyses and the candidate genes set from the studies outlined above.

## Supplementary Material

Supplementary materials are available for this manuscript.

## Acknowledgements

This research utilised Queen Mary’s Apocrita HPC facility, supported by QMUL Research-IT (http://doi.org/10.5281/zenodo.438045). We thank past and present members of the Plant Health and Adaptation Team at RBG Kew for helpful discussions and advice on various aspects of this research. We thank D. Wood for useful comments on an earlier version of this manuscript. This work was supported by the Living with Environmental Change Tree Health and Plant Biosecurity Initiative – Phase 2, funded jointly by BBSRC, Defra, ESRC, Forestry Commission, NERC and the Scottish Government [(grant no. BB/L012162/1]). L.J.K. acknowledges additional support from RBG Kew.

## Data Availability

The data underlying this article are available in its online supplementary material and in Zenodo at doi.org/10.5281/zenodo.20412204 and doi.org/10.5281/zenodo.2041000; additional original data can be accessed at the European Nucleotide Archive with accession number PRJEB20151 and in Zenodo at doi.org/10.5281/zenodo.6948951.

## Notes

https://zenodo.org/records/20412204

### Competing Interest Statement

The authors have declared no competing interest.

### Summary of Updates

Additional text has been added to the Discussion and Materials and Methods sections, and minor changes made to other sections of the text to improve clarity. None of the results or figures have been changed.

