## Supplementary Figures for "Convergent molecular changes are associated with the evolution of low susceptibility to the ash dieback pathogen"

### Molecular convergence analyses identify candidate genes for low susceptibility to the ash dieback pathogen

#### Supplementary Figures

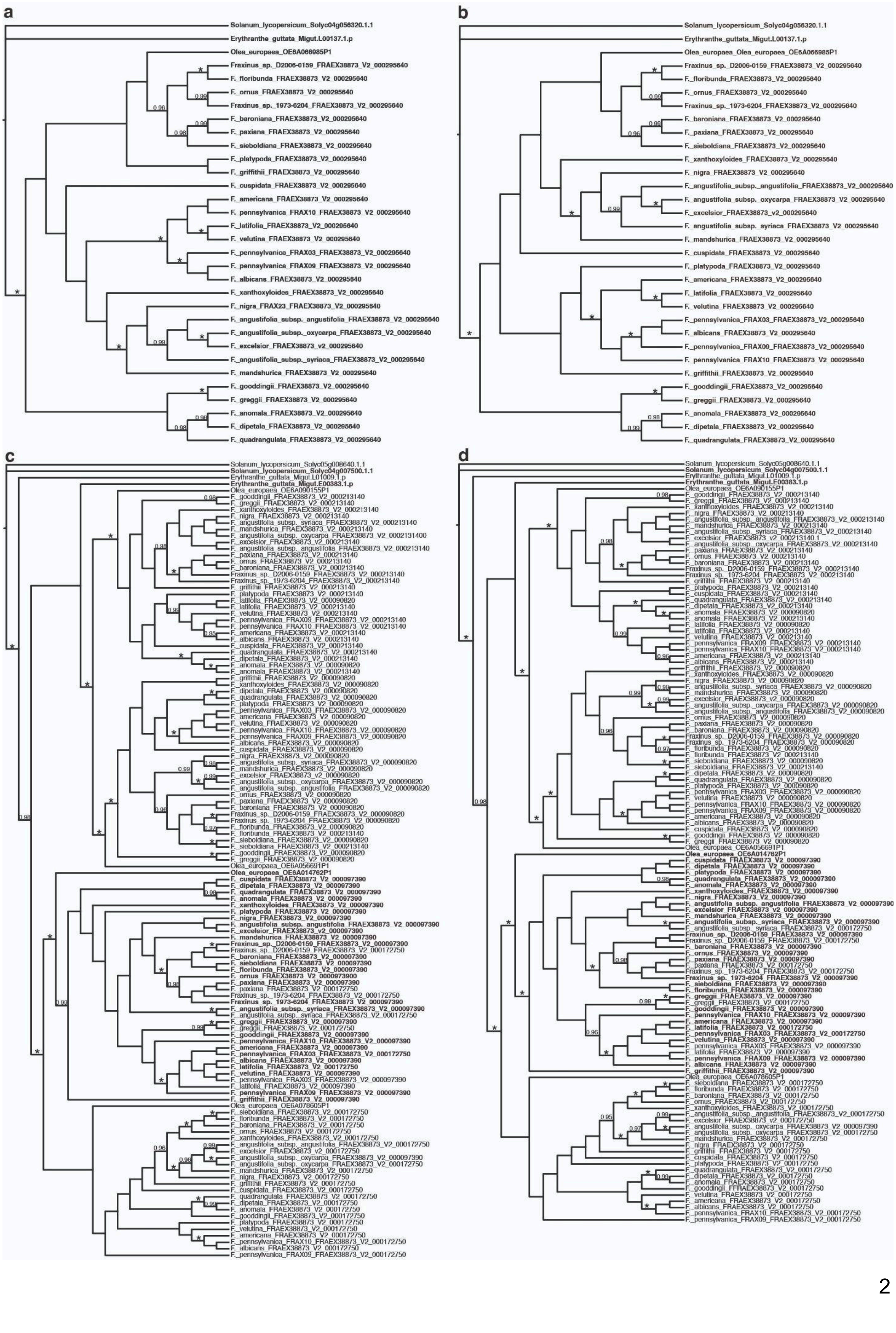
a
Solanum_lycopersicum_Solyc04g056320.1.1
Erythranthe_guttata_Migut.L00137.1.p
Olea_europaea_OE6A066985P1
Fraxinus_sp._D2006-0159_FRAEX38873_V2_000295640
F._floribunda_FRAEX38873_V2_000295640
F._ornus_FRAEX38873_V2_000295640
Fraxinus_sp._1973-6204_FRAEX38873_V2_000295640
F._baroniana_FRAEX38873_V2_000295640
F._paxiana_FRAEX38873_V2_000295640
F._sieboldiana_FRAEX38873_V2_000295640
F._platypoda_FRAEX38873_V2_000295640
F._griffithii_FRAEX38873_V2_000295640
F._cuspidata_FRAEX38873_V2_000295640
F._americana_FRAEX38873_V2_000295640
F._pennsylvanica_FRAX10_FRAEX38873_V2_000295640
F._latifolia_FRAEX38873_V2_000295640
F._velutina_FRAEX38873_V2_000295640
F._pennsylvanica_FRAX03_FRAEX38873_V2_000295640
F._pennsylvanica_FRAX09_FRAEX38873_V2_000295640
F._albicans_FRAEX38873_V2_000295640
F._xanthoxyloides_FRAEX38873_V2_000295640
F._nigra_FRAX23_FRAEX38873_V2_000295640
F._angustifolia_subsp._angustifolia_FRAEX38873_V2_000295640
F._angustifolia_subsp._oxycarpa_FRAEX38873_V2_000295640
F._excelsior_FRAEX38873_v2_000295640
F._angustifolia_subsp._syriaca_FRAEX38873_V2_000295640
F._mandshurica_FRAEX38873_V2_000295640
F._gooddingii_FRAEX38873_V2_000295640
F._greggii_FRAEX38873_V2_000295640
F._anomala_FRAEX38873_V2_000295640
F._dipetala_FRAEX38873_V2_000295640
F._quadrangulata_FRAEX38873_V2_000295640
b
Solanum_lycopersicum_Solyc04g056320.1.1
Erythranthe_guttata_Migut.L00137.1.p
Olea_europaea_Olea_europaea_OE6A066985P1
Fraxinus_sp._D2006-0159_FRAEX38873_V2_000295640
F._floribunda_FRAEX38873_V2_000295640
F._ornus_FRAEX38873_V2_000295640
Fraxinus_sp._1973-6204_FRAEX38873_V2_000295640
F._baroniana_FRAEX38873_V2_000295640
F._paxiana_FRAEX38873_V2_000295640
F._sieboldiana_FRAEX38873_V2_000295640
F._xanthoxyloides_FRAEX38873_V2_000295640
F._nigra_FRAEX38873_V2_000295640
F._angustifolia_subsp._angustifolia_FRAEX38873_V2_000295640
F._angustifolia_subsp._oxycarpa_FRAEX38873_V2_000295640
F._excelsior_FRAEX38873_v2_000295640
F._angustifolia_subsp._syriaca_FRAEX38873_V2_000295640
F._mandshurica_FRAEX38873_V2_000295640
F._cuspidata_FRAEX38873_V2_000295640
F._platypoda_FRAEX38873_V2_000295640
F._americana_FRAEX38873_V2_000295640
F._latifolia_FRAEX38873_V2_000295640
F._velutina_FRAEX38873_V2_000295640
F._pennsylvanica_FRAX03_FRAEX38873_V2_000295640
F._albicans_FRAEX38873_V2_000295640
F._pennsylvanica_FRAX09_FRAEX38873_V2_000295640
F._pennsylvanica_FRAX10_FRAEX38873_V2_000295640
F._griffithii_FRAEX38873_V2_000295640
F._gooddingii_FRAEX38873_V2_000295640
F._greggii_FRAEX38873_V2_000295640
F._anomala_FRAEX38873_V2_000295640
F._dipetala_FRAEX38873_V2_000295640
F._quadrangulata_FRAEX38873_V2_000295640
c
d

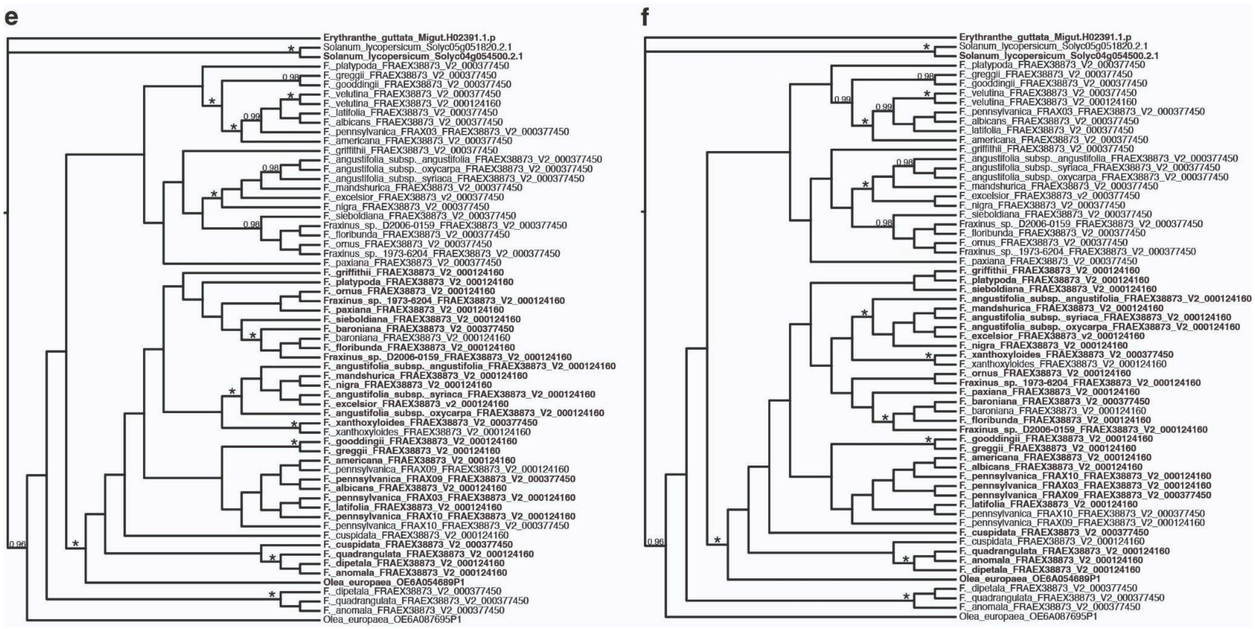


Supplementary Fig. 1. Gene-trees for hierarchical orthologous groups (HOGs) where the sequences including convergent amino acid residues grouped together more closely than expected from the species-tree. a) Gene-tree for HOG2851 (which encompasses OG27879, where convergence was detected between *F. platypoda* and members of section *Ornus*). b) Gene-tree for HOG2851 with the position at which convergence was detected excluded. c) Gene-tree for HOG614 (which encompasses OG61784, where convergence was detected between *F. mandshurica* and members of section *Ornus*). d) Gene-tree for HOG614 with the position at which convergence was detected excluded. e) Gene-tree for HOG3654 (which encompasses OG64792, where convergence was detected between *F. platypoda* and members of section *Ornus*). f) Gene-tree for HOG3654 with the position at which convergence was detected excluded. Tip labels for sequences found in the OGs are shown in bold; note that for HOG2851 the sequence membership is identical to that of its equivalent OG and thus all tip labels are bold. Gene-trees were inferred with MrBayes from protein multiple sequence alignments with unreliably aligned positions removed. Numbers above branches are posterior probabilities (PP) of ≥0.95; asterisks indicate nodes with PP=1. Gene model names follow species names; for *F. pennsylvanica* (which is represented by data from three different individuals) genome sequencing codes used in Kelly et al. (2020) precede the gene model names.

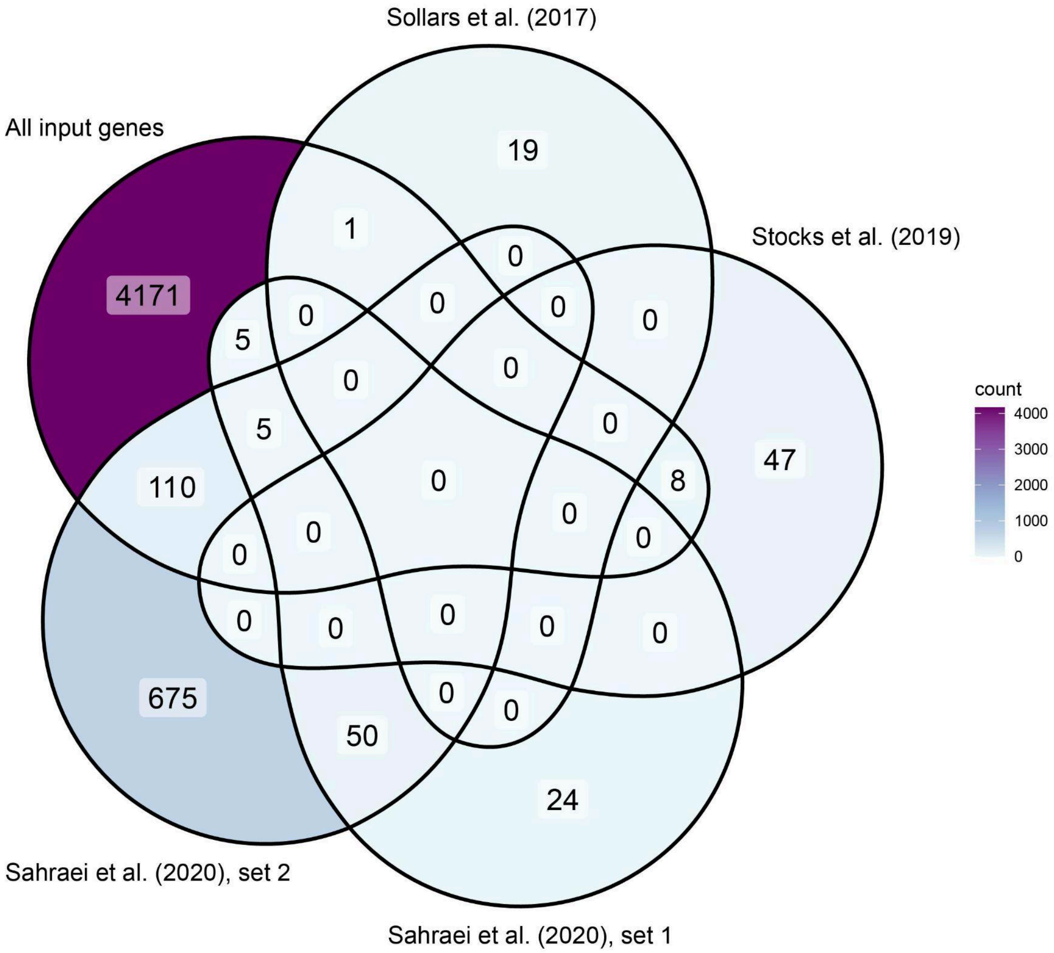


Supplementary Fig. 2. Venn diagram showing overlap between the set of 4,300 loci used as input for the molecular convergence analyses and those associated with level of susceptibility to *H. fraxineus* in *F. excelsior* identified from previous analyses. Gene sets identified in previous studies are: Sollars et al.(2017) - 20 gene expression markers (GEMs) identified via associative transcriptomics; Stocks et al.(2019) - 55 genes containing or within 5 kb up or downstream of a SNP significantly associated with health status in a GWAS; Sahraei et al.(2020), set 1 - 84 genes with evidence of differential expression between inoculated *F. excelsior* individuals with varying levels of susceptibility, identified from sequencing of symptomatic tissue; Sahraei et al. (2020), set 2 - 840 genes with evidence of differential expression between inoculated *F. excelsior* individuals with varying levels of susceptibility, identified from sequencing of asymptomatic tissue.
